# Quantifying the impacts of an invasive weed on habitat quality and prey availability for tiger snakes (*Notechis scutatus*) in urban wetlands

**DOI:** 10.1101/2022.12.07.519536

**Authors:** Jari Cornelis, Brenton von Takach, Christine E. Cooper, Jordan Vos, Philip. W. Bateman, Damian C. Lettoof

**Affiliations:** School of Molecular and Life Sciences, Curtin University, Western Australia, Australia; Independent Researcher, Secret Harbour 6173, Western Australia, Australia

**Keywords:** Elapidae, Kikuyu grass, Urbanisation, Predation, Environmental temperature, Habitat degradation

## Abstract

Invasive plants are a threat to natural ecosystems worldwide with urban wetlands being some of the most susceptible and highly modified environments of all. The tiger snake (*Notechis scutatus*) is a top predator that persists in urban wetlands in south-western Australia, many of which have been degraded by introduced kikuyu grass (*Cenchrus clandestinus*). To evaluate the potential impact of kikuyu grass on habitat quality for western tiger snakes we quantified the structural features of habitats within wetlands degraded by kikuyu grass and compared them to wetlands with native vegetation, examined tiger snake prey availability, assessed predation risk for juvenile snakes using clay models, and measured the thermal quality of the vegetation. Proliferation of kikuyu grass reduced habitat structural heterogeneity by reducing available bare ground and increasing vegetation density. This homogenisation of habitat structure had little effect on the predation risk for juveniles or the thermal properties of tiger snake shelter sites; however, one key snake prey species, the motorbike frog, had significantly lower abundance in the most impacted habitat. Habitat types with more structural complexity also offered tiger snakes more stable thermal regimes and lower predation risk. These findings indicate that the current extent of kikuyu grass invasion offers overall similar habitat quality for tiger snakes and may contribute to their persistence in urban wetlands, but they, along with their anuran prey, may befit from some increased habitat structural complexity in open areas.

## Introduction

Urbanisation is a major driver of global environmental change, with urban infrastructure and human populations encroaching on and modifying natural ecosystems (Faulkner 2004, Coleman 2017, Cresswell and Murphy 2017). Urbanisation leads to the degradation and removal of natural spaces for plants and animals and the introduction of invasive of species which create new biological interactions and alter the physical environment (McDonnell et al. 1993, Kinzig and Grove 2001, Hamer and McDonnell 2008, Štajerová et al. 2017). Invasive plants are a threat to ecosystems worldwide and can dominate disturbed or sensitive vegetation communities. This can reduce plant diversity and structural heterogeneity, impacting ecosystem function (Schweiger et al. 2018) and retention of biodiversity (Stein et al. 2014). Monocultures of weeds can reduce habitat quality for fauna by reducing structural complexity and resource availability, impacting predation pressure, thermal quality, food availability and ultimately the overall suitability of the habitat (Huey 1991, Webb et al. 2005, Lee et al. 2006).

Large reptiles are the top predators of many Australian ecosystems, with diverse effects on ecosystem structure and function (Pianka 1986, Kuch et al. 2005, Read and Scoleri 2015, Doody et al. 2021). Habitat use and selection by snakes is influenced by its thermal properties, with ideal habitats offering a selection of thermally optimal microhabitats (Hertz et al. 1994) balancing thermoregulation opportunities, prey abundance and shelter from predation (Amo et al. 2007). Tiger snakes (*Notechis scutatus*), which grow to about 1 m long, are a viviparous species of the elapid (Elapidae) family and occur throughout southern and eastern Australia. Tiger snakes are typically affiliated with wetlands and predominately prey on amphibians, but considerable behavioural and morphological plasticity allows them to persist in a diverse array of habitats (Shine 1987, Aubret and Shine 2010) including urban areas (Butler et al. 2005). In the south-west of the continent, western tiger snakes (*N. s. occidentalis*) are a top predator of wetland ecosystems (Aubret 2005), with their presence and abundance sensitive to an intact food web and habitat availability (Sergio et al. 2008). Western tiger snakes persist in a subset of urban wetlands within the urban sprawl of Perth city (Lettoof et al. 2021c) where they are considered bio-indicators of wetland health (Lettoof et al. 2021b) and prey predominantly on frogs (Shine 1998, Lettoof et al. 2020).

Some of the urban wetlands inhabited by western tiger snakes are dominated by introduced kikuyu grass (*Cenchrus clandestinus*), a globally recognised invasive species, particularly in wetlands (Bird et al. 2013, Boon and Tesfamichael 2017). Abundant kikuyu lowers native plant species richness and cover (Gaertner et al. 2011) and has potentially similar impacts on fauna. However, not all fauna is negatively affected by the invasion of exotic grasses (Douglas et al. 2006). For some reptiles habitat structure is more important than plant species composition (Garden et al. 2007) and populations may persist in highly modified habitats if certain structural components are retained within the landscape. For example, the presence of structurally complex habitats created by the invasive woody shrub lantana (*Lantana camara*) supports large numbers of a rare species of shadeskink (*Saproscincus rosei;* Virkki et al. 2012). Invasive grasses such as kikuyu may also provide suitable habitat for the frogs on which tiger snakes prey (Zavaleta et al. 2001, Maerz et al. 2005) and prey availability is an important driver of snake abundance and habitat use (McCauley et al. 2006, Battles et al. 2013, Zipkin et al. 2020).

A broadly inhospitable matrix of urban area likely restricts tiger snakes in Perth to the wetlands they inhabit (Lettoof et al. 2021c), with little opportunity for individuals to move between habitat patches. Consequently, the quality of these wetlands and availability of prey are factors crucial for maintenance of viable population sizes. An over-abundance of kikuyu appears to have substantially changed the vegetation composition and habitat structure of these wetlands. This could have flow-on effects for prey abundance, predation risk and thermal quality for tiger snakes, but this has not yet been studied. Here, we investigate the impacts of kikuyu invasion on these variables, by comparing a range of biotic and abiotic metrics in kikuyu-dominated wetlands against wetlands that are dominated by native vegetation communities. We quantify vegetation structure, prey availability (frog abundance), predation risk (for juveniles), and thermal quality of available microhabitats to better understand how kikuyu might impact this urban predator and assist land managers to mitigate the degradation of urban ecosystems and their biodiversity.

## Methods

### Study sites

We examined western tiger snake habitats at two wetlands dominated by kikuyu and two wetlands dominated by native vegetation, all within 50 km of the Perth CBD, in south-west Western Australia (Fig 1). Western tiger snake populations have been previously documented as occurring at these wetlands (Lettoof et al. 2021c). At Herdsman Lake (HL; 31.92°S, 115.80°E) and Kogolup Lake (KL; 32.12°S, 115.83°E) kikuyu was the dominant ground cover vegetation while at Loch McNess in Yanchep National Park (Y; 31.54°S, 115.68°E) and Black Swan Lake (BS; 32.47°S, 115.77°E) native vegetation predominated. Within these four sites, we identified three *a priori* broad habitat categories; open grassland (abundant ground cover of either uncut kikuyu grass, or native grasses with no canopy cover or mid-story vegetation), sedge (mid-story level dense native sedges or uncut kikuyu grass growing against bullrush bordering waterbodies), and woodland habitats (canopy cover and understory vegetation, occasionally with mid-story vegetation; Fig S1) and use the terms ‘kikuyu’ or ‘native’ to distinguish the broad wetland communities from the finer-scale habitat categories within these sites.

**Fig. 1.**
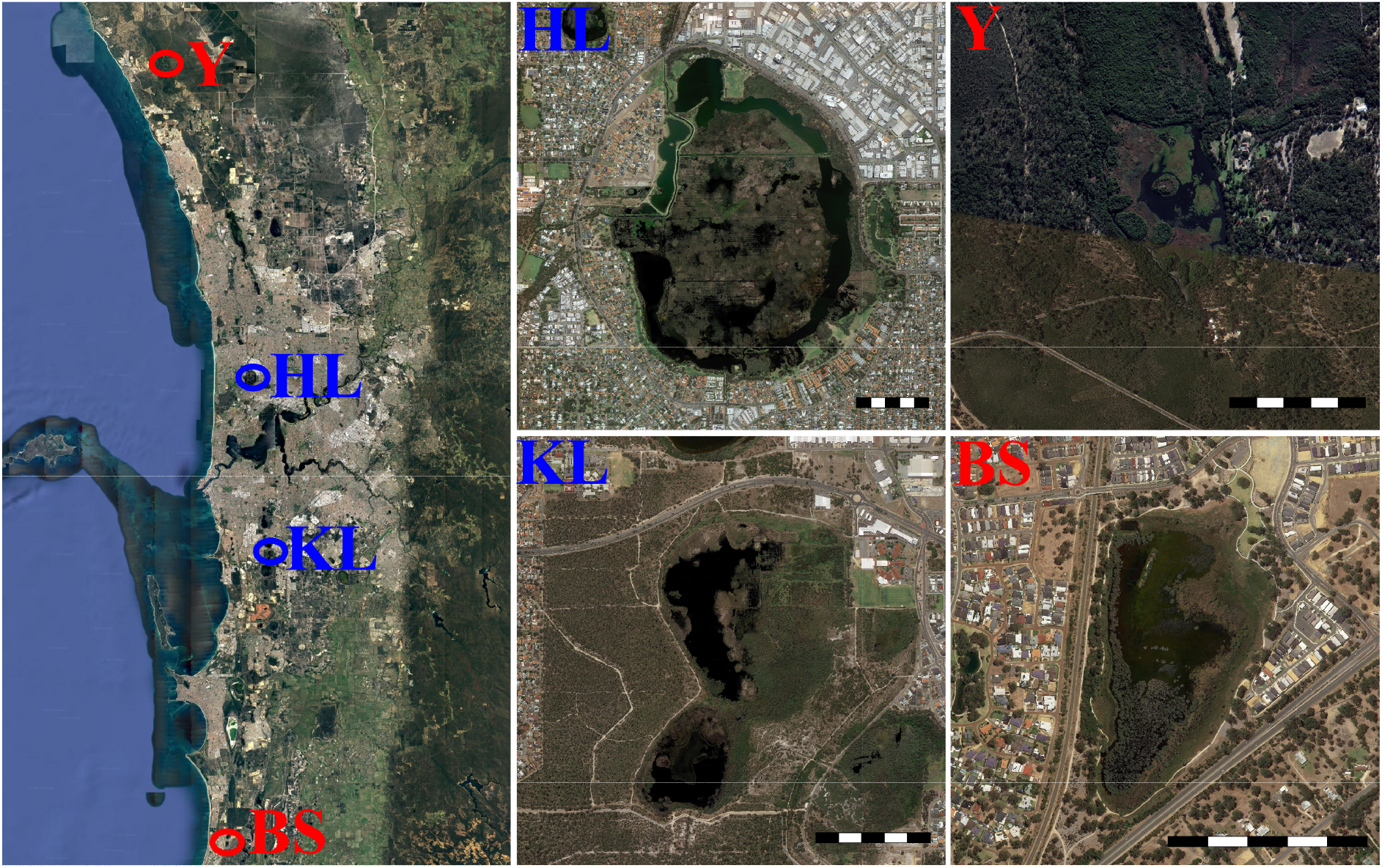
Wetlands that were surveyed in the Swan Coastal Plain. BS = Black Swan Lake, HL = Herdsman Lake, KL = Kogolup Lake, Y = Yanchep National Park. Red font indicates wetlands classed as dominated by native vegetation, blue font indicates wetlands dominated by kikuyu grass. Satellite images were obtained from Google Earth in 2022. Scale bar = 500m.

### Habitat structure

To quantify habitat structure, we recorded vegetation features along two 100m transects in each habitat type at each wetland. Every 10m along the transect we recorded canopy cover using a spherical densiometer (Forest Densiometers, Model A), light availability with a digital lux meter (DR.meter LX101BS, when the reading in ambient unshaded light was ca. 720,000 lux), the number of trees with stems >150mm circumference at breast height (1.4m) within a 2m radius, the number of woody plants with stems <150mm circumference at breast height within a 2m radius, the height of the tallest understorey plant, an estimate of the percent ground cover within a 2m radius, and the vegetation density. Vegetation density was quantified by inserting a 50 cm x 50 cm board inside the vegetation (2 m from the observer), taking a photograph, and then using ImageJ 1.x (Schneider et al. 2012) to calculate the surface area of the board not obscured by vegetation. If the vegetation was too dense to insert the board we recorded vegetation density as 100% (Fox et al. 1996, Colman et al. 2014, Colman et al. 2015).

All statistical analyses were conducted in R studio version 1.4.1103 (RStudio Team 2020). The lme4 package (Bates et al. 2014) was used for all generalised linear mixed-effects models, with the emmeans package (Lenth et al. 2018) used to conduct Tukey’s post-hoc pairwise comparisons. We used principal component analysis (PCA) to examine associations between the three *a priori* habitat types and vegetation communities (i.e. kikuyu or native vegetation) based on the measured habitat characteristics. The FactoMineR (Lê et al. 2008) and factoextra (Kassambara and Mundt 2020) packages were used to generate principal components (PCs); PCs with eigenvalues > 1 were considered useful for inference (Roznik and Reichling 2021). An analysis of deviance was then used to compare the PCs of the habitat types and to compare sites with kikuyu to those with native vegetation, with Tukey’s *post-hoc* pairwise comparisons used to determine differences between habitat characteristics (Roznik and Reichling 2021).

### Prey availability

We measured the abundance of prey (frogs) in the different vegetation communities by conducting standardised 10 minute active visual searches in the open, sedge and woodland habitat types at the four wetlands. The search commenced in the centre of each habitat type and observers walked a steady pace while counting individual frogs detected with a spotlight. Frogs were identified to species. Only a single wetland was surveyed each night to mitigate any temporal effects caused by time of night. Sites Y and BS (native) were surveyed for five nights each, and KL and HL (kikuyu) were surveyed on four nights each, for a total of 54 visual searches. Surveys were conducted in October and November (2019), which is peak calling season for the dominant prey species *Litoria moorei, Litoria adelaidensis* and *Limnodynastes dorsalis* (Hoskin et al. 2015, Lettoof et al. 2020). All surveys began after 19:00 on evenings when ambient temperature at this time was >15°C and were completed before 22:30.

Generalised linear mixed-effects models (GLMMs) were used to assess the differences in frog abundance among vegetation communities and habitats. We used a two-step process for assessing differences: Firstly, frog abundance was broadly compared between vegetation communities and habitat types; fitting a GLMM (Poisson error structure) with frog count data as the response variable, vegetation community (kikuyu or native vegetation) and habitat type as fixed effects and site as a random effect. The second approach was to compare frog abundance to habitat structure variables to determine which specific structural components of a habitat type were being selected, aiding in assessing the fine-scale effects of kikuyu vegetation changes for the target frog species; fitting a GLMM (Poisson error structure) with frog count data as the response variable, the habitat structure variables as predictor variables, and site as a random effect. Predictor variables were first scaled and centred to improve model fitting, and correlated variables were removed to reduce multicollinearity, retaining variables with variance inflation factors < 5 and pairwise correlations < 0.7, identified using the usdm and psych packages (Naimi 2015, Revelle 2015, Lettoof et al. 2020). For correlated variables the least ecologically relevant predictors were removed (Dormann et al. 2013). The global model included canopy cover, vegetation density, vegetation height and woody vegetation as predictor variables. Model fit was then assessed with pseudo-R^2^ values calculated with the glmmADMB package (Skaug et al. 2013). Following this, model selection was performed using the dredge function from the MuMIn package (Barton 2015) and all sub models were ranked according to Akaike’s information criterion, with a sample size correction (AICc). All models with ΔAICc < 2 were considered useful for inference.

### Predation risk

We assessed the differences in predation risk for young tiger snakes among vegetation communities and habitats using clay models. We moulded 120 soft clay models, representing juvenile tiger snakes because snakes are most vulnerable to predation when they are small (Lima and Dill 1990, Webb et al. 2005), and 120 spherical control balls from nontoxic Plastiplay^TM^ brown modelling clay (Fig S2A). Total body length of the artificial snakes was ca. 25cm, width 1cm, and they were moulded into a sinusoidal shape. Control balls were 4cm in diameter and used to determine if predators discriminated foreign objects from artificial snakes (Nordberg and Schwarzkopf 2019b). The artificial snakes and balls were deployed at the four study sites in each of the three different habitat types. In each habitat type, ten artificial snakes were placed in exposed areas to simulate basking sites, and ten were hidden under vegetation to simulate refuge sites. Artificial snakes were separated by 5m, with each model accompanied by a control ball at a distance of 25cm. All clay models were checked every day for a five day period and marks left by predators were counted, scored by intensity (depth) of attack, and the predators were identified as birds, rodents or cats, based on the shape of the marks left by the predators (Webb and Whiting 2005), or classified as unknown if the clay models were missing. Small shallow marks, often left by house mice (*Mus musculus*), were scored as exploratory rather than predatory as they did not represent a realistic predatory attempt on a juvenile tiger snake; these were therefore omitted from analyses (Fig S2B). Experiments were conducted in autumn and repeated in spring for a total of 10 days of sampling at each site.

We used a Student’s t-test to compare the difference in predation rate between snake models and control balls. We used the same two-step process used to analyse frog abundance data for assessing differences in predation counts both among vegetation communities and habitat types, and for habitat structure variables. The response variable for this GLMM was whether a model had experienced a predation attack (binomial yes/no). This global model included vegetation density, ground cover, light availability and vegetation height as predictor variables.

### Thermal quality of vegetation

To determine if kikuyu grass or native vegetation differ in their thermal quality we monitored the temperatures within potential tiger snake shelter sites with operative temperature models constructed using 30cm long sections of copper pipe (32mm diameter, wall 1mm thick) filled with water and painted to approximate the reflectivity of a tiger snake (Seebacher and Shine 2004, Lutterschmidt and Reinert 2012). In the centre of each model we suspended a small temperature logger (Thermochron iButton) which recorded temperature at 10min intervals. The temperature logger was waterproofed (Plasti Dip) and wrapped in plastic film. Temperatures of the operative models were calibrated against tiger snake carcasses over a range of thermal and radiative conditions (Fig S3). We deployed models in the field and calculated their thermal properties via a custom-written VB program (Visual Basic V6; Cooper and Withers 2004).

Three operative temperature models were deployed at each wetland, one in each of the three habitat types, placed within what we refer to as a shelter site (deep within the vegetation where we have observed tiger snakes retreating or emerging; pers obs.). Data were recorded at 10min intervals for seven days every month for twelve months. We calculated the mean, maximum and minimum operative environmental temperature (T_e_) of the sheltered microhabitats. Then we calculated the thermal quality (*de*) of tiger snake shelter sites as the absolute value of how much T_e_ in shelter sites deviated from the thermal set-point range (T_set_) for tiger snakes (Blouin-Demers and Weatherhead. 2002), determined by Ladyman & Bradshaw (2003) as 24.0 ± 0.9°C to 30.9 ± 0.3°C for snakes from HL in a thermal gradient. The resulting data were then grouped into four seasons and we compared the differences in thermal quality between sites with kikuyu grass and native vegetation, and the habitat types within sites. To compare seasonal habitat temperature characteristics between vegetation types, we used a GLMM (Gaussian error structure) with the temperature characteristic as the response variable, and vegetation type, habitat type and season as fixed predictor variables, with site as a random effect. Tukey’s post-hoc pairwise comparisons were used to examine differences between variables.

## Results

### Habitat structure

Three PCs were retained (eigenvalue >1) from the PCA analysis of habitat structure, collectively accounting for 73.7% of the total variance. PC1 explained 35.5% of the variance, and was associated with canopy cover, number of trees, woody vegetation and low light availability (Table S1). PC2 explained 23.8% of the variance and was associated with ground cover, vegetation density and canopy cover. PC3 explained 14.4% of the variation and was associated with understory height. Collectively, the structure of kikuyu grass habitats was more homogenous than native vegetation habitats (Fig 2) and differed significantly for PC2 (F_5, 238_ = 495, P < 0.001). The structure of our *a priori* habitat types were significantly different for PC1 (F_5, 234_ ≤ 149, P < 0.001), although no significant differences were detected for kikuyu open (KO) and native open (NO; P = 0.526), or for kikuyu sedge (KS) and native sedge (NS; P = 0.989), or kikuyu woodland (KW) and native woodland (NW; P = 0.998) in PC2. For PC3, KO and NO were statistically similar (P = 0.856) while KS and NS, and KW and NW differed (P < 0.001).

**Fig. 2.**
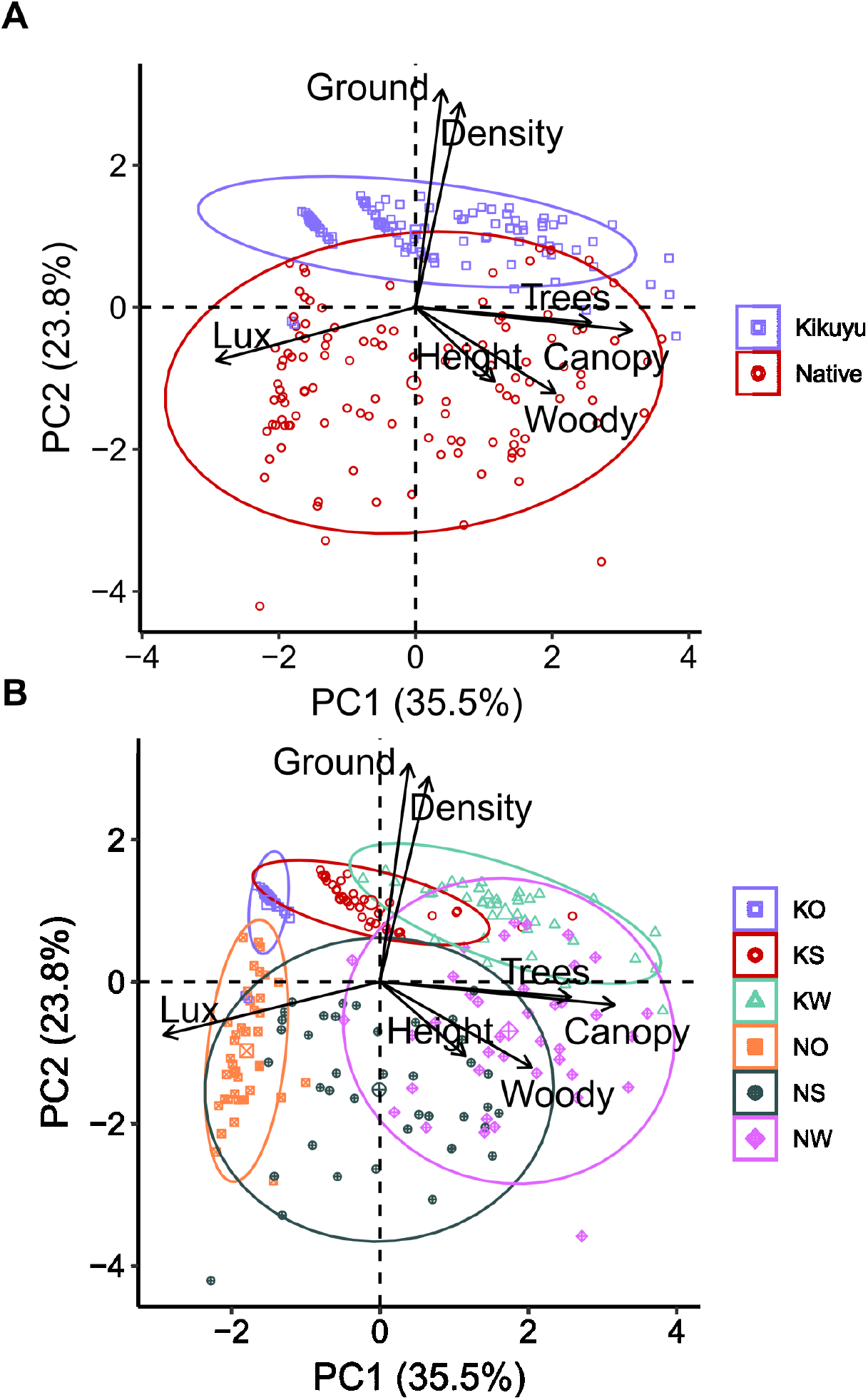
PCA ordination of habitat characteristics in western tiger snake (*Notechis scutatus occidentalis*) habitats from four different wetlands, dominated by either invasive vegetation (Herdsman Lake and Kogolop Lake) or native vegetation (Black Swan Lake and Yanchep National Park) in the Perth region, Western Australia. (a) the ordination of the habitat characteristics of the kikuyu grass and native vegetation plant communities (b) ordination of the different habitat types within the wetlands KO, kikuyu open; KS, kikuyu sedge; KW, kikuyu woodland; NO, native open; NS, native sedge; NW native woodland and. PC1 explains 35.5% and PC2 21% of the total variance. Arrows denote the influence of specified habitat characteristics and ellipses denote 80% of the spread of the groups.

### Prey availability

Three species of frog were frequently observed during surveys: slender tree frog (*Litoria adelaidensis*), moaning frog (*Heleioporus eyrie*) and motorbike frog (*Litoria moorei*). Two individuals of the rattling froglet (*Crinia glauerti*) were detected at a single site (Yanchep) and the moaning frog was not detected at Herdsman Lake (Table 1). Consequently, the rattling froglet was excluded from all subsequent analyses and the moaning frog was included in the ‘total frogs’ analysis (but with insufficient sample size to explore habitat preferences).

**Table 1.** Number of frogs seen during visual surveys at four wetlands in Perth, Western Australia. Wetlands include Herdsman Lake (HL), Kogolup Lake (KL), Black Swan Lake (BS) and Yanchep National Park (Y). Frog species are the slender tree frog (*Litoria adelaidensis;* STF), moaning frog (*Heleioporous eyrei;* MF) and motorbike frog (*Litoria moorei;* MBF).

|  |  | STF | MF | MBF |
| --- | --- | --- | --- | --- |
| Kikuyu | HL | 14 | 0 | 56 |
|  | KL | 38 | 9 | 78 |
| Native | BS | 227 | 2 | 114 |
|  | Y | 51 | 11 | 95 |

Total frog abundance was not significantly influenced by vegetation community (kikuyu grass or native vegetation; χ^2^_1_ = 3.19, P = 0.076); however, the habitat types (open, sedge and woodland) and the interaction between vegetation community and habitat types did influence the total abundance of frogs (χ^2^_2_ ≤ 34.23, P ≤ 0.004), with a greater abundance of frogs in the native open (NO) habitat than in the kikuyu open habitat (KO; *post-hoc* comparison P = 0.007; Fig 3). Further, there were significant differences in the abundance of frogs among habitat types within vegetation communities. Native open habitats had significantly more frogs than the native sedge (NS) and native woodland habitats (NW; *post-hoc* comparisons P < 0.001). In kikuyu grass communities, frog abundance was significantly lower in the open habitat compared to the kikuyu woodland (KW; *post-hoc* comparisons P = 0.001).

**Fig. 3.**
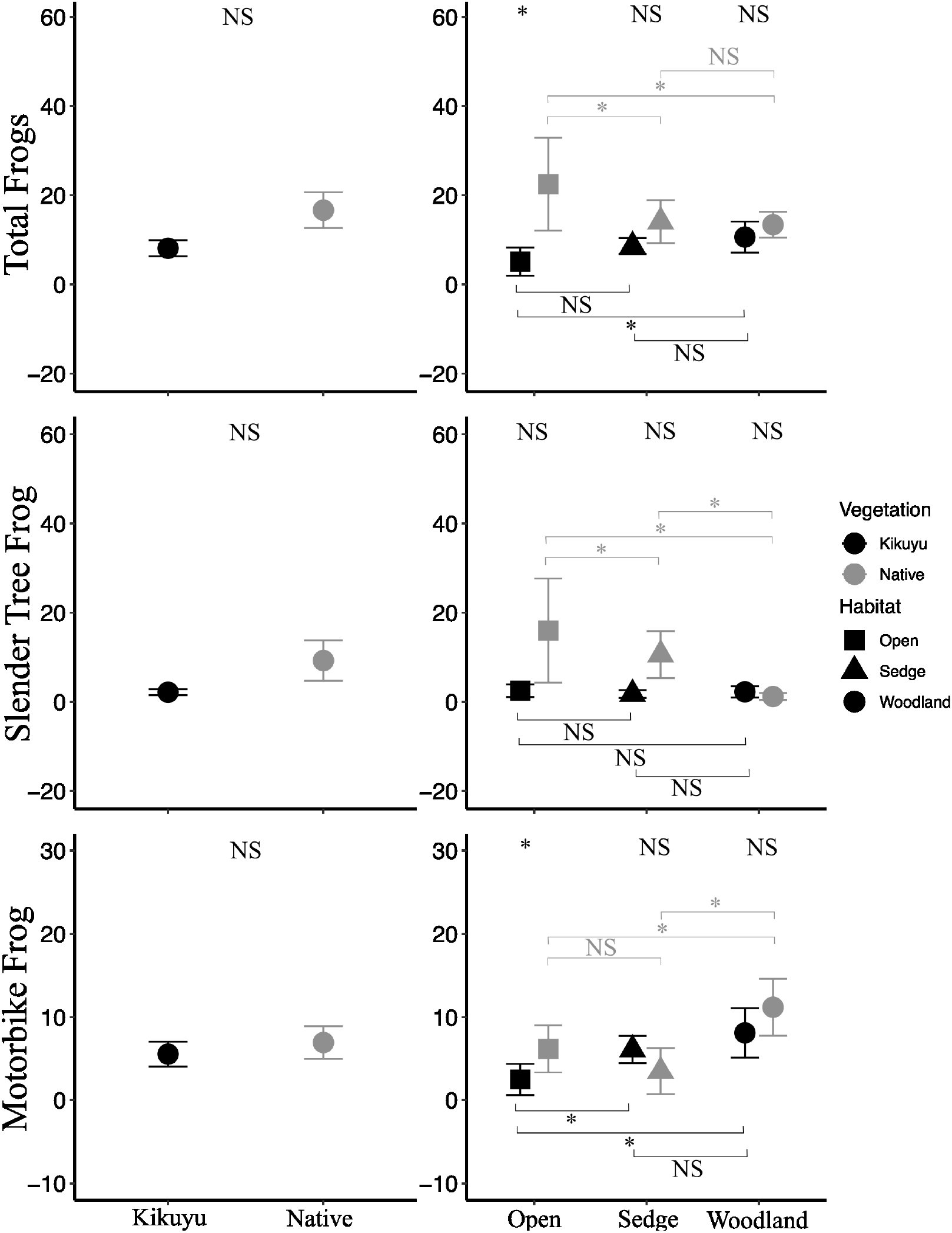
Mean (± SE) frog abundance in the Perth region, Australia, in wetlands dominated by either kikuyu grass or native vegetation. Panels a) and b) show patterns in total frogs (all species summed), panels c) and d) show patterns for slender tree frogs (*Litoria adelaidensis*), and panels e) and f) show patterns for motorbike frogs (*Litoria moorei*). * denotes a significant difference between habitat types within vegetation communities.

The abundance of the slender tree frog (χ^2^_1_ = 3.54, P = 0.059) and motorbike frog (χ^2^_1_ = 2.53, P = 0.111) did not differ between vegetation communities (Fig 3). However habitat types, and the interaction between the habitat types and vegetation communities influenced the abundance of both species (slender tree frog: χ^2^_2_ ≤ 44.1, P < 0.001; motorbike frog: χ^2^_2_ ≤ 41.68, P < 0.001). The slender tree frog and motorbike frog were similar in abundance between direct equivalent habitat types i.e. KO and NO (*post-hoc* comparisons; P ≥ 0.072), except the motorbike frog was significantly more abundant in NO than KO (*post-hoc* comparisons P = 0.004). Within kikuyu grass communities, slender tree frog abundance was not different (*post-hoc* comparisons P ≥ 0.820), but the motorbike frog was significantly more abundant in woodlands (KW) than in open and sedge habitats (KO and KS; *post-hoc* comparisons P ≤ 0.006). Within native vegetation communities, slender tree frog abundance was significantly different among all habitat types (*post-hoc* comparisons P ≤ 0.008), and the motorbike frog was more abundant in woodlands than open and sedge habitats (*post-hoc* comparisons P ≤ 0.001). Canopy cover, vegetation density and ground cover (%) were important predictors of both species; slender tree frog abundance was negatively associated with canopy cover and ground cover, and positively associated with vegetation density (Table 2), whereas motorbike frog abundance was positively associated with canopy cover, ground cover and understorey height, and as was negatively associated with vegetation density (Table 2).

**Table 2.**
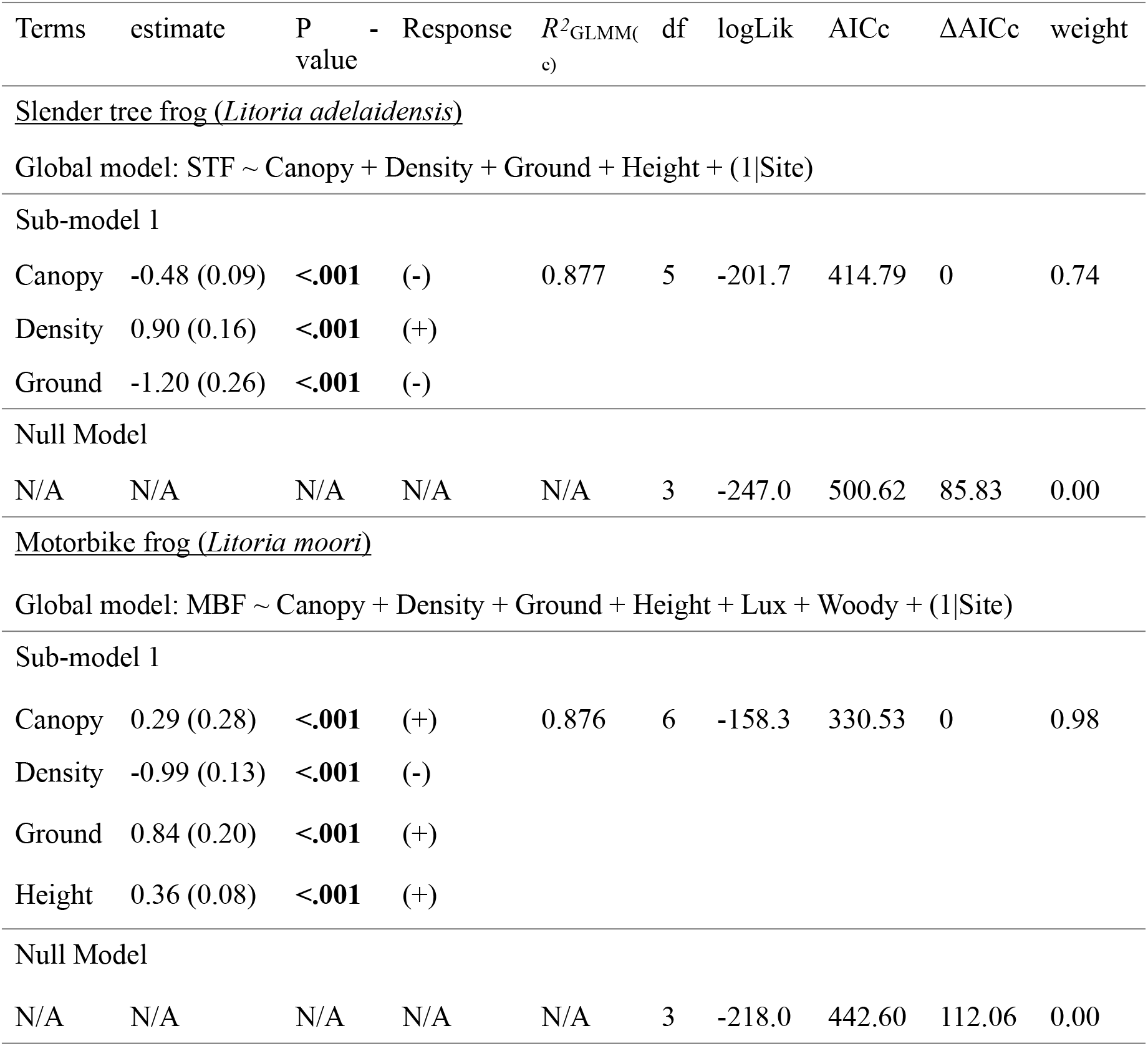
Habitat characteristics influencing the abundance of slender tree (*Litoria adelaidensis*) and motorbike frogs (*Litoria moori*). Results from generalised linear mixed-effects model indicating the best predictor variables for submodels with ΔAICc < 2 and the null model identifying the strongest predictors of abundance based on ΔAICc and weight in four wetlands in the Perth region, Western Australia. The estimate and standard error (estimate) and P-value are reported for each predictor variable. The conditional R^2^ (*R*_GLMM(c)_), degrees of freedom (df), log-likelihood (logLik), Akaike’s Information Criterion (AICc), difference in AICc between models (ΔAICc), and weight are reported for each model. Response indicates whether frog abundance had a positive (+) or negative (-) response to the habitat characteristics. Significant p-values are in bold.

### Predation risk

Of the 1200 observations of clay models, 57 predation attempts were recorded for artificial snakes and 17 for control balls. Artificial snakes were significantly more likely to be predated than control balls (t958 = 3.98, P < 0.001). Most predation marks could be identified as being a consequence of either birds (54%) or rodents (40%; Table 3). We could not investigate differences in the frequency of different predator interactions with the models due to the infrequency of predation attempts. All predation data were therefore pooled into a single response variable, ‘predation’.

**Table 3.** Predation attempts on artificial snakes in vegetation communities that are either dominated by either introduced kikuyu grass or native plant species. Predation attempts were identified as being made by a bird, rodent, cat, or unknown species, and were quantified in three habitat types (classified as open, sedge, and woodland).

|  |  | Bird | Rodent | Cat | Unknown |
| --- | --- | --- | --- | --- | --- |
| Kikuyu | Open | 10 | 1 | 0 | 0 |
|  | Sedge | 9 | 10 | 0 | 1 |
|  | Woodland | 6 | 0 | 0 | 0 |
| Native | Open | 4 | 3 | 1 | 0 |
|  | Sedge | 2 | 7 | 0 | 0 |
|  | Woodland | 0 | 2 | 0 | 1 |

There was no significant difference in the probability of predation on artificial snakes between vegetation communities (χ^2^_1_ = 1.27, P = 0.257). The probability of predation was significantly different among habitat types (χ^2^_2_ = 6.86, P = 0.032) although t *post-hoc* comparisons of pairwise habitats showed no significant differences (P ≥ 0.270; Fig 4). Light availability and woody vegetation were the best predictors of predation probability for artificial snakes, as these variables occurred in most of the five top models (Table 4). Predation probability increased with light availability and decreased with woody vegetation count.

**Fig. 4.**
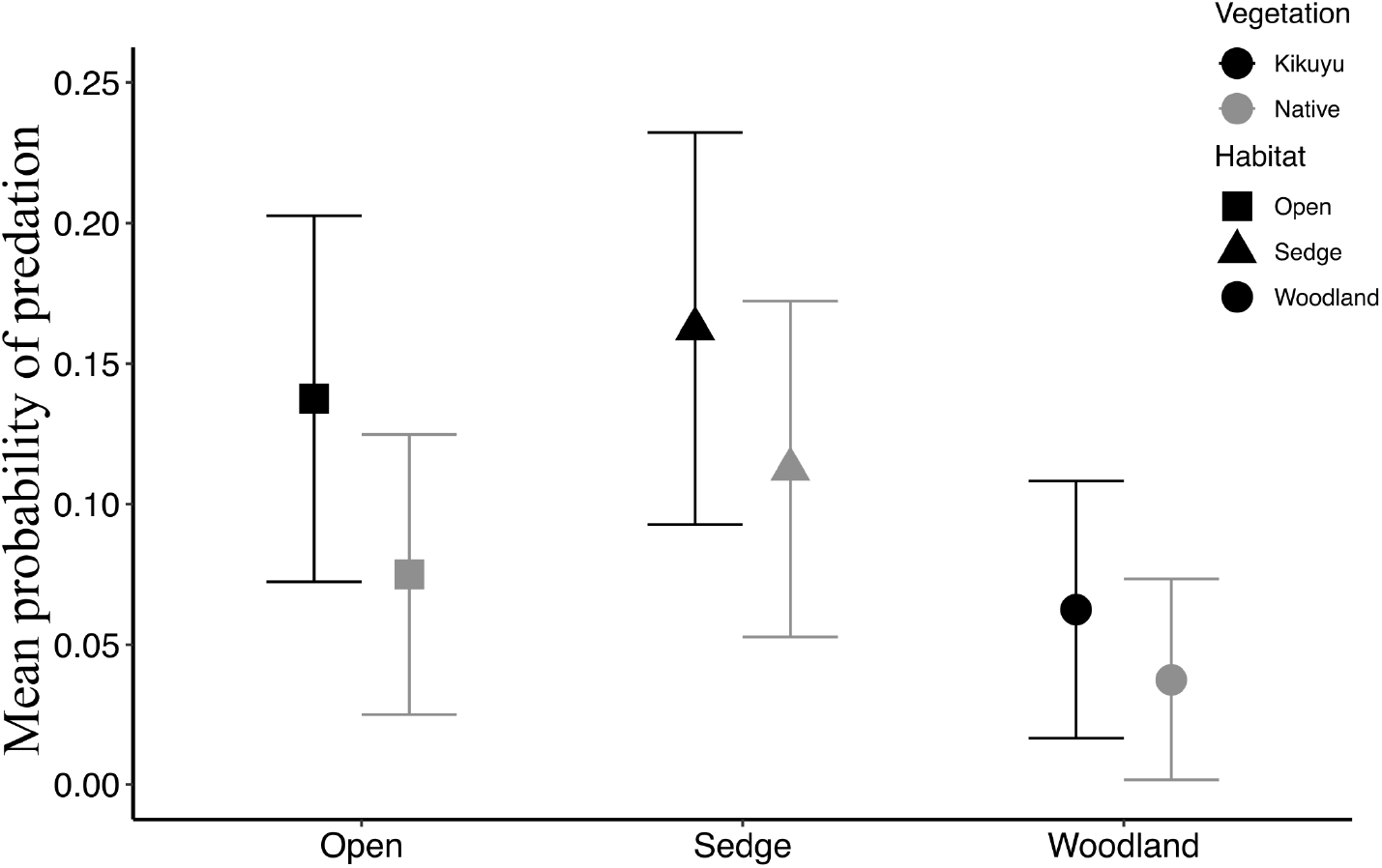
Mean (± SE) probability of predation for clay model snakes in wetland vegetation communities dominated by either introduced kikuyu grass (*Cenchrus clandestinus*) or native vegetation in Perth, Western Australia. Predation attempts were quantified in three habitat types (open, sedge and woodland).

**Table 4.** Habitat characteristics influencing the predation risk for clay models of juvenile snakes. Submodels and the null model identifying the strongest predictor variables of clay model snake predation probability based on ΔAICc < 2 and weight in four wetlands in the Perth region, Western Australia. The estimate and standard error (estimate) and P-value are reported for each predictor variable. The conditional *R^2^* (*R*_GLMM(c)_), degrees of freedom (df), log-likelihood (logLik), Akaike’s Information Criterion (AICc), difference in AICc between models (ΔAICc), and weight are reported for each model. Response indicates whether predation probability had a positive (+) or negative (-) response to the habitat characteristics. Significant p-values are in bold.

| Terms | estimate | P-value | Respons<br>e | $R^2_{GLMM(c)}$ | df | logLik | AICc | $\Delta AICc$ | weight |
| --- | --- | --- | --- | --- | --- | --- | --- | --- | --- |
| Global model: Attacks ~ Density + Ground + Height + Lux + Woody + (1\|Site) |  |  |  |  |  |  |  |  |  |
| Sub-model 1 |  |  |  |  |  |  |  |  |  |
| Lux | 0.28 (0.14) | **0.049** | (+) | 0.126 | 4 | -147.5 | 303.18 | 0 | 0.19 |
| Woody | -0.46 (0.27) | 0.090 | (-) |  |  |  |  |  |  |
| Sub-model 2 |  |  |  |  |  |  |  |  |  |
| Woody | -0.55 (0.27) | **0.047** | (-) | 0.110 | 3 | -149.4 | 304.87 | 1.69 | 0.08 |
| Sub-model 3 |  |  |  |  |  |  |  |  |  |
| Lux | 0.34 (0.14) | **0.016** | (+) | 0.097 | 3 | -149.4 | 304.89 | 1.71 | 0.08 |
| Sub-model 4 |  |  |  |  |  |  |  |  |  |
| Height | 0.09 (0.16) | 0.573 | (+) | 0.130 | 5 | -147.3 | 304.91 | 1.73 | 0.08 |
| Lux | 0.30 (0.14) | **0.043** | (+) |  |  |  |  |  |  |
| Woody | -0.47 (0.27) | 0.084 | (-) |  |  |  |  |  |  |
| Sub-model 5 |  |  |  |  |  |  |  |  |  |
| Density | -0.04 (0.20) | 0.820 | (-) | 0.130 | 5 | -147.5 | 305.17 | 1.99 | 0.07 |
| Lux | 0.28 (0.14) | 0.051 | (+) |  |  |  |  |  |  |
| Woody | -0.45 (0.27) | 0.095 | (-) |  |  |  |  |  |  |
| Null Model |  |  |  |  |  |  |  |  |  |
| N/A | N/A | N/A | N/A | N/A | 5 | -151.9 | 313.95 | 10.77 | 0.00 |

### Thermal quality of vegetation

Calibration of the operative temperature models indicated they accurately represented the temperatures an actual tiger snake would experience (see description in Online Resource 1 and Fig S3). Most of the thermal variables were highly correlated with *de_max_* (R ≥ 0.90), except for T_e, max_ (R = 0.68) and *de_0_* (proportion of time *de* was equal to 0; R = 0.43), so we only analysed vegetation and habitat effects for these three variables. The T_e,max_ differed seasonally (F_3_ = 413.63, P = < 0.001), being highest in summer and lowest in winter (Fig 4). The T_e,max_ in habitat types did not differ within each season (*post-hoc* comparisons P ≥ 0.190), besides the native open habitat which was significantly higher than the native sedge during summer (*post-hoc* comparisons P < 0.001; Fig 5). Vegetation community had no influence on *de_max_* (F_1_ = 0.05, P = 0.837). There was a significant difference in *de_max_* among habitat types (F_2_ = 31.79, P < 0.001) with the highest *de_max_*s occurring in open habitats and the lowest *de_max_*s in the woodland habitats (Fig 5). Season also significantly influenced *de_max_* (F_3_ = 275.53, P < 0.001) which was lowest during summer, highest during winter and similar in autumn and spring (Fig 5), with no interaction (F_6_ = 0.21, P = 0.972; Fig 5). Neither vegetation community (F_1_ = 0.04, P = 0.857) or habitat type (F_2_ = 1.69, P = 0.184) influenced the mean proportion of time that the thermal quality of the vegetation fell within T_set_ (*de_0_*). However there was a significant interaction between vegetation community, habitat type and season (F_6_ = 3.04, P = 0.005); within a season, there were no differences between equivalent habitat types in different vegetation communities (*post-hoc* comparisons P ≥ 0.240) but there were some differences between habitat types within the same vegetation community e.g. the highest mean *de_0_* occurred in KW during summer which was different from KO and KS in the same season (*post-hoc* comparisons P ≤ 0.001; Fig 5).

**Fig. 5.**
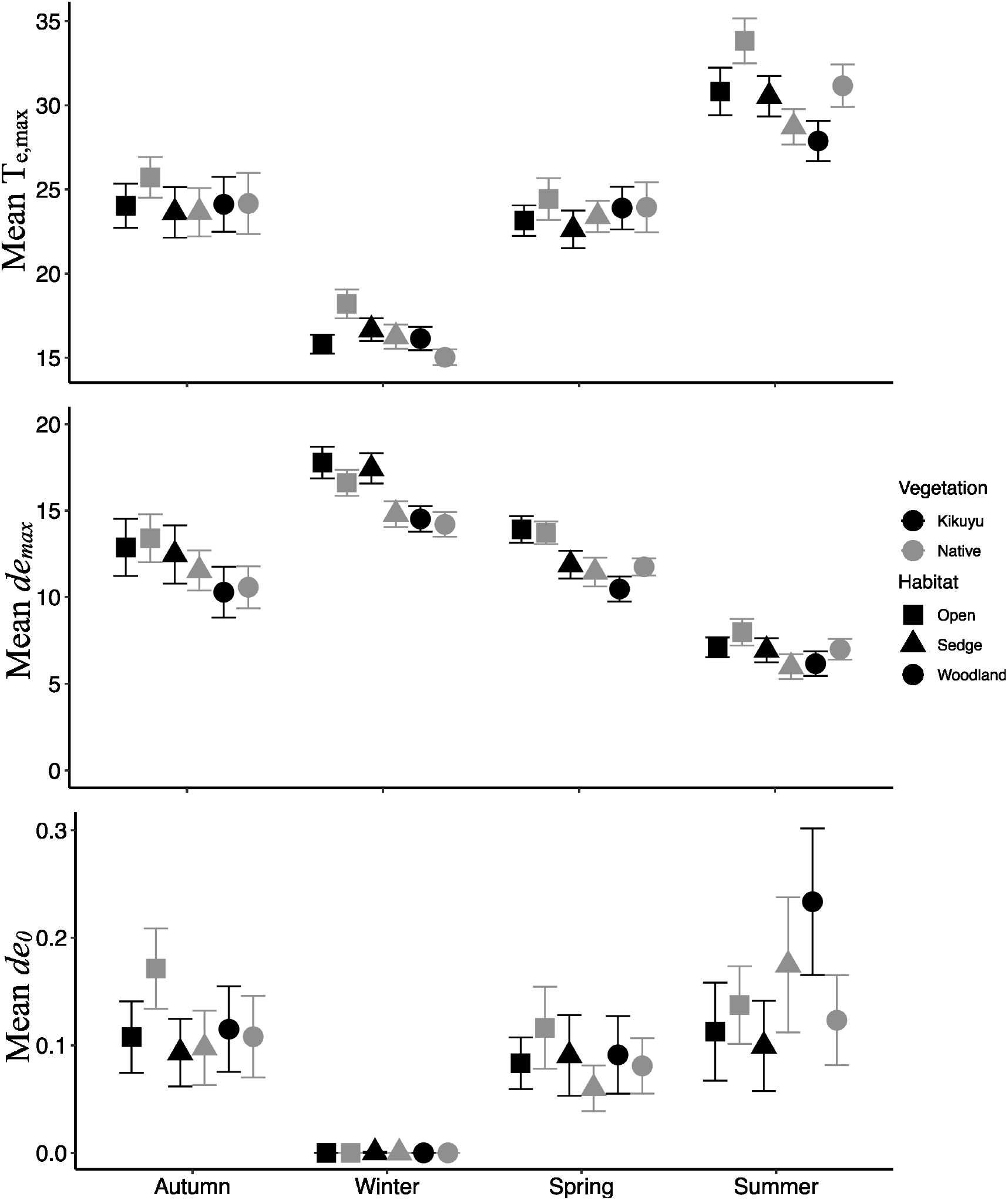
Temperature parameters (A; maximum operative environmental temperature; T_e_, B; maximum deviation of environmental temperature, *de_max_* and C; proportion of time the environmental temperature in sheltered microhabitats fell within the body temperature set-point range; *de_0_*) for tiger snakes (*Notechis scutatus occidentalis*) at four wetlands in Perth, Western Australia recorded during four seasons. Parameters were measured in sheltered microhabitats in three habitat types at sites where the vegetation is dominated by either introduced kikuyu grass or native vegetation.

## Discussion

We investigated a suite of habitat factors likely to influence populations of western tiger snakes in urban wetlands where native vegetation communities have been largely replaced by a single invasive species, kikuyu grass. Our results indicate that habitat compositional heterogeneity at sites with kikuyu grass has been reduced, and that prey availability may be reduced in areas that have become dominated by kikuyu. However, despite the reduction in bare ground and increase in vegetation density at sites invaded by kikuyu, the transition from native vegetation to invasive grass appears to have had little effect on the predation risk for juvenile snakes or the thermal properties of shelter sites. Together, these findings indicate that the current extent of invasion of kikuyu grass offers an overall similar habitat quality for tiger snakes to natural vegetation and is unlikely to substantially impact population persistence in urban wetlands.

### Habitat structure

Diverse habitat structure is a primary driver of biodiversity worldwide (Culbert et al. 2013, Boeye et al. 2014, von Takach et al. 2020). Our results indicated that habitat structure is different between kikuyu-dominated and native-dominated vegetation, but is similar between habitat types (open/sedge/woodland), supporting our *a priori* determination of habitat categories. Increased vegetation density, ground cover and overall homogenisation drove the observed differences in the structure of kikuyu dominated vegetation communities and native vegetation equivalents. Kikuyu grows in a dense matrix of stems, which facilitates its colonisation and reduces inter-plant distance compared to native grasses, resulting in less bare ground and reduced plant diversity, transforming native vegetation communities into a monoculture (Gonzalez 2009, Litt and Steidl 2011, Lindsay and Cunningham 2012, Bradshaw et al. 2013, Abom et al. 2015). The domination of an environment by a monoculture of a single invasive grass can have adverse effects on the ecosystem. For example, the invasion of gamba (*Andropogon gayanus*) and para (*Urochloa mutica*) grasses in northern Australia results in higher fuel loads and higher intensity fires than in uninvaded habitats, due to higher biomass and lower moisture content of the invasive species (Rossiter et al. 2003, Douglas and O’Connor 2004, von Takach et al. 2022). The most common impact of habitat structural homogeneity from invasive grasses, however, is the loss of native plant (Stephens et al. 2008), invertebrate (Douglas and O’Connor 2003) and vertebrate (Cook and Grice 2013, Stanton et al. 2018) biodiversity – largely attributed to a loss of suitable habitat usually provided by structurally complex vegetation communities. Similar to these studies, we found that the invasion of kikuyu grass changed the habitat structure of riparian wetland vegetation. While we quantified the impact of this change on two common frog species, we encourage further research into how this invasion may affect wetland biota.

### Prey availability

In general, current levels of kikuyu invasion do not impact total abundance of frogs. However, the abundance of slender tree frog and motorbike frog among particular habitat types differed within and between these vegetation communities. The slender tree frog was found in higher abundance in the native vegetation communities of all three habitat types, with the lowest abundance found in open kikuyu habitats. Slender tree frogs are small (34-47mm) prey for tiger snakes, so a lack of slender tree frogs in kikuyu-dominated habitats likely reduces prey availability for young tiger snakes. The motorbike frog seemed more tolerant of kikuyu grass invasion, but had the highest abundance in woodland habitats and a negative association with vegetation density. Motorbike frogs contribute to a substantial proportion of tiger snake diet in these wetlands (Lettoof et al. 2020), and their persistence—and contribution to tiger snake diet biomass—may be reliant on riparian woodland habitat.

The open kikuyu habitats apparently had the lowest abundance of all frogs, although it is possible, due to the dense structure created by kikuyu grass, we were unable to detect all frogs sheltering in kikuyu grass (Heard et al. 2008, Vences et al. 2008). This habitat type is a monoculture of dense grass, and structurally very different from the open native vegetation equivalent which is comprised of several structurally diverse plant species and interspersed with bare ground. This is unsurprising, as invasive grass monocultures usually result in a loss of abundance and diversity of native birds (Catling 2005, Skórka et al. 2010), reptiles (Valentine et al. 2007, Hacking et al. 2014), frogs (Grant and Samways. 2016, Falaschi et al. 2020) and rodents (Sammon and Wilkins. 2005) globally. The abundance of these two critical prey species was highest in sedge and woodland habitats where the greater structural complexity and woody vegetation is likely favourable (Landsman and Bowman 2017), potentially reducing predation risk (Norbury and van Overmeire 2019). As large expanses of wetland riparian vegetation in Perth can be kikuyu grass monocultures (Hill et al. 1996, Department of Environment and Conservation 2012), the persistence of these frogs and their tiger snake predators could be facilitated by the planting and maintenance of riparian sedge (*Gahnia decomposita* and/or Lepidosperma longitudinale) and native woodland trees (Banksia, Melaleuca and Eucalyptus spp).

### Predation risk

Predation attempts on juvenile snake clay models occurred at low frequency, with birds and rodents the predominant predators. The low frequency of predation attempts (4.75% of 1200) on artificial juvenile tiger snakes is comparable to the predation rates (7.44%) observed for artificial juvenile broad headed snakes (*Hoplocephalus bungaroides*; Webb and Whiting 2006) and uropeltid snakes (5%; Cyriac and Kodandaramaiah 2019). Using clay models as a proxy for live animal interactions can have limitations (Bateman et al. 2017), yet we consider our predation attempts to be an accurate representation of at least bird and rodent predation at our study sites, as predation attempts were significantly higher on snake models compared to control balls. The depredated model snakes had strike marks primarily on the head and tail and occasionally had been flipped by an avian predator and pecked on the ventral surface, typical predation strategies for predators of snakes (Webb and Whiting 2006). It is likely that predation of juvenile tiger snakes at these sites is in reality a rare occurrence and indeed the observed predation rate could be an overestimate as live juvenile snakes spend limited time basking (Webb and Whiting 2005) and artificial snakes do not respond with antipredator behaviours.

Environmental habitat structure influences the predation risk for reptiles by reducing the protective qualities of refuge sites or increasing structural features predators use to hunt (Hawlena et al. 2010, Martin and Murray 2011, Steidl et al. 2013). The dominant predators at our study sites are largely visual hunters, thus the artificial snakes were probably more conspicuous in habitats that have reduced structural complexity and increased light availability (Daly et al. 2008, Sato et al. 2014). This is consistent with other studies that found greater predator diversity along open habitat edges (Anderson and Burgin 2008). Birds specifically use elevated perches along habitat edges to forage in open environments (Hansen et al. 2019) and preference hunting in more homogenous habitats (Hawlena et al. 2010). Our results suggest that kikuyu dominated vegetation is not influencing predation risk for juvenile tiger snakes but riparian vegetation with woody stemmed vegetation and structural complexity provides safer habitats for young tiger snakes, highlighting the value of maintaining sedge and woodland habitats in these urban wetlands.

### Thermal quality of vegetation

Kikuyu grass dominated habitats had similar thermal properties to those dominated by natural vegetation. Similarly, Abom et al. (2015) found that the thermal quality within grader grass (*Themeda quadrivalvis*) did not differ from within native vegetation, offering thermally equivalent shelter sites for a number of skink and snake species. As the thermal environments within these vegetation communities are so similar, despite a change in habitat composition, the way tiger snakes thermoregulate and exploit the thermoregulatory opportunities in these environments has remained unaffected (Cornelis 2021). Not all vertebrate species are disadvantaged by the invasion of exotic grasses (Malo et al. 2012, Lindenmayer et al. 2017); the thermal quality and dense structure provided by kikuyu grass may be particularly suitable for snakes as they can easily move through the vegetation and thermoregulate (McDonald and Luck 2013, Abom et al. 2015). The only notable differences were that woodland habitats provided the most stable thermal environments with the least extreme temperature fluctuations, which is unsurprising given the effects of canopy cover in providing shade and reducing radiation to the sky (Breshears et al. 1998, Yates et al. 2000). In summer in kikuyu dominated communities, woodlands provided the highest proportion of optimal tiger snake temperature compared to other habitats (Table S2). The most challenging temperatures for tiger snakes occur in summer and winter. In Perth during summer, Ta regularly exceeds 40^°^C and winter nights can drop below 0°C (Bureau of Meteorology 2022). Snakes are at considerable risk of exceeding their voluntary thermal maxima (35.5°C) in summer or critical thermal minima (2°C) during winter (Shine 1977, Lillywhite 1980, Shine and Mason 2004). In some cases, habitat types in either vegetation community approached these temperatures but the thermal stability, with generally lower maximum and higher minimum temperatures, offered by the woodland habitat types may offer tiger snakes the most valuable thermoregulatory and hibernation opportunities (Table S2).

### Management and conservation implications

Seventy percent of urban wetland habitat on the Swan Coastal Plain between Wedge Island and Mandurah Western Australia has been lost to urban development and agriculture (Halse 1989). Of the remaining 30%, only 17% is vegetated with predominantly native vegetation (Hill et al. 1996). Weeds have been identified as a key threat to Perth metropolitan wetlands (Conservation and Parks Commission 2017) and as such weed management and rehabilitation plans have been implemented by many local governments and/or the Department of Biodiversity, Conservation, and Attractions. Complete eradication of kikuyu grass and revegetation with native species can be extremely difficult (Department of Environment and Conservation 2012) and may not be successful in some situations. Our work suggests that total eradication is not necessary to maintain western tiger snake populations in wetlands of the Swan Coastal Plain and large-scale removal of kikuyu may leave snakes and their prey without shelter sites. We suggest that an alternative to eradication that may improve habitat quality for the snakes and their major prey items would be to increase structural habitat complexity, particularly in areas of homogenous open kikuyu habitat. This could be achieved by removing smaller plots of kikuyu grass and planting more structurally complex native species, with scattered canopy cover. A suitably large area of kikuyu grass would need to be removed to allow for the establishment of these slower growing native species as perennial grasses such are kikuyu are known to quickly smother native plants (Department of Environment and Conservation 2012). Though this method may result in positive outcomes for local western tiger snake and frog populations, more research is needed to determine how kikuyu dominated vegetation is used by other fauna, and the potential impacts of restoration on other flora and fauna species within the wetland ecosystem.

In Perth, tiger snakes are only known to persist in seven of the potential hundreds of available wetlands (Hill et al. 1996, Lettoof et al. 2021a). Although the exact habitat requirements necessary for supporting tiger snake populations are unknown, we suspect adequate vegetation structure plays a vital role. Over-grown kikuyu grass provides the necessary structural features in wetlands where native riparian vegetation no longer exists, providing suitable prey, anti-predator and thermal functions. Future research efforts should focus on assessing which habitat characteristics allow tiger snakes to persist in urban wetlands, how tiger snakes utilise the habitats we described in this study (e.g. space-use, movement and habitat selection preferences), and the population size and stability of remnant populations.

## Acknowledgments

This study was supported by a grant from the Holsworth Wildlife Research Endowment to DCL. The authors thank Aleesha Turner, Serin Subaraj and Brae Price for their assistance in the field. We also thank the staff at the Department of Biodiversity Conservation and Attractions and the staff at Regional Parks for allowing us access to the study sites and issuing the required permits.

## Declarations

### Ethical approval

Experiments followed the Australian Code of Practice for the care and use of animals for scientific purposes and were approved by the Curtin University Animal Ethics committee (ARE2019-24).

### Author contributions

All authors contributed to the study conception and design. Material preparation, data collection and analysis were performed by Jari Cornelis, Brenton von Takach and Damian Lettoof. The first draft of the manuscript was written by Jari Cornelis and all authors commented on previous versions of the manuscript. All authors read and approved the final manuscript. Additional funding was provided by Damian Lettoof.

### Data availability

All relevant raw data and R scripts are available on request from the authors.

