## Supplementary Material for "Quantifying the impacts of an invasive weed on habitat quality and prey availability for tiger snakes (*Notechis scutatus*) in urban wetlands"

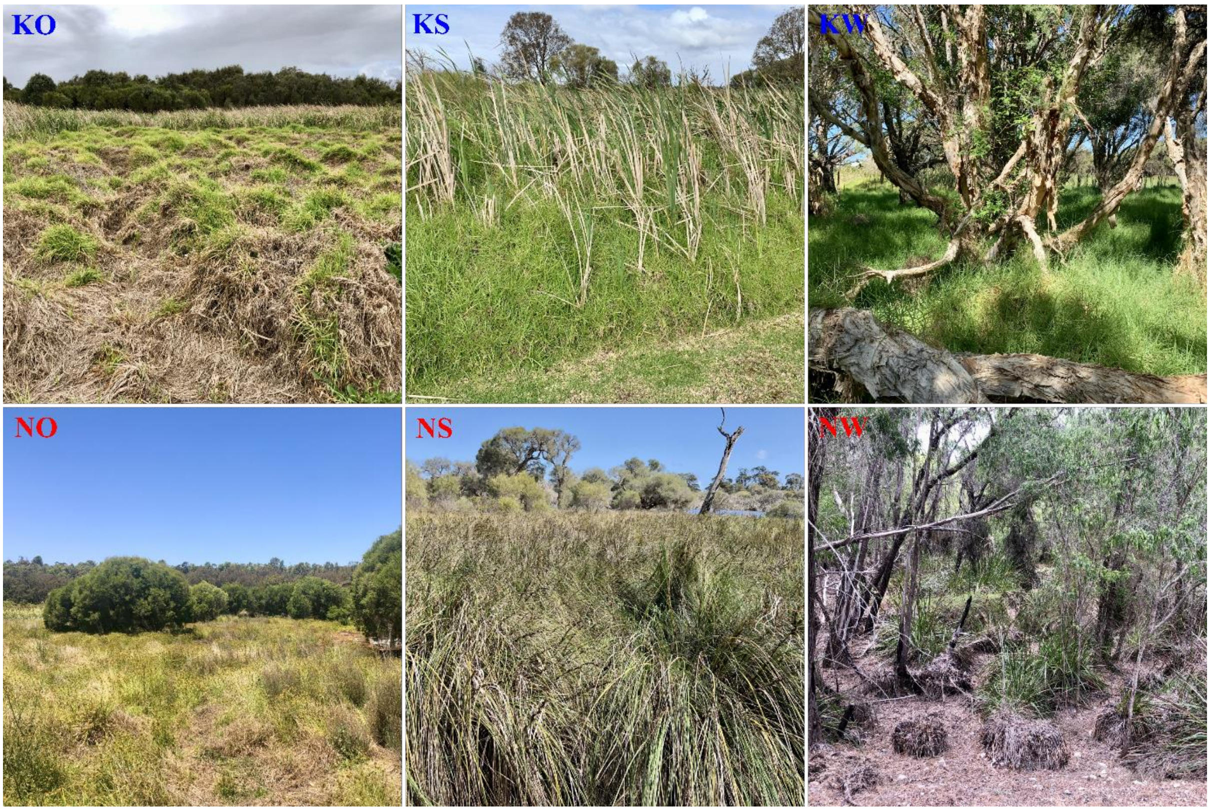


**Fig S1** Habitat types surveyed at four wetlands on the swan coastal plain. The open kikuyu habitat (KO) was a dense mat of kikuyu without canopy cover, the kikuyu sedge habitat (KS) was categorised by a dense, tall (up to 2m) mass of kikuyu growing on a secondary structure provided by bulrush (*Typha* sp.) and the kikuyu woodland habitat (KW) had a dense mat of kikuyu growing in the shade of *Eucalyptus*, *Melaleuca* and/or *Banksia spp.*. The native open grassland habitat (NO) was a mixed community with the dominant grass type *Schoenoplectus spp.* The native sedge habitat (NS) consisted of large tussock species: *Gahnia decomposita* and/or *Lepidosperma longitudinale*; while the native woodland habitat (NW) was a mixed understory community growing beneath *Banksia*, *Melaleuca* and *Eucalyptus spp.*

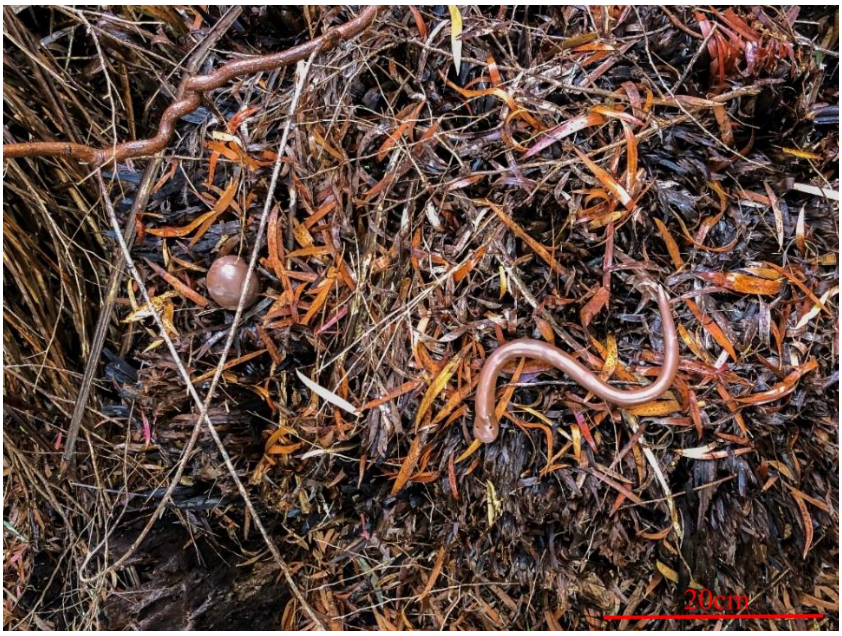


**Fig S2A** Artificial snake and control ball made from nontoxic Plastiplay$^{TM}$ brown modelling clay used to quantify predation risk for juvenile western tiger snakes (*Notechis scutatus occidentalis*).

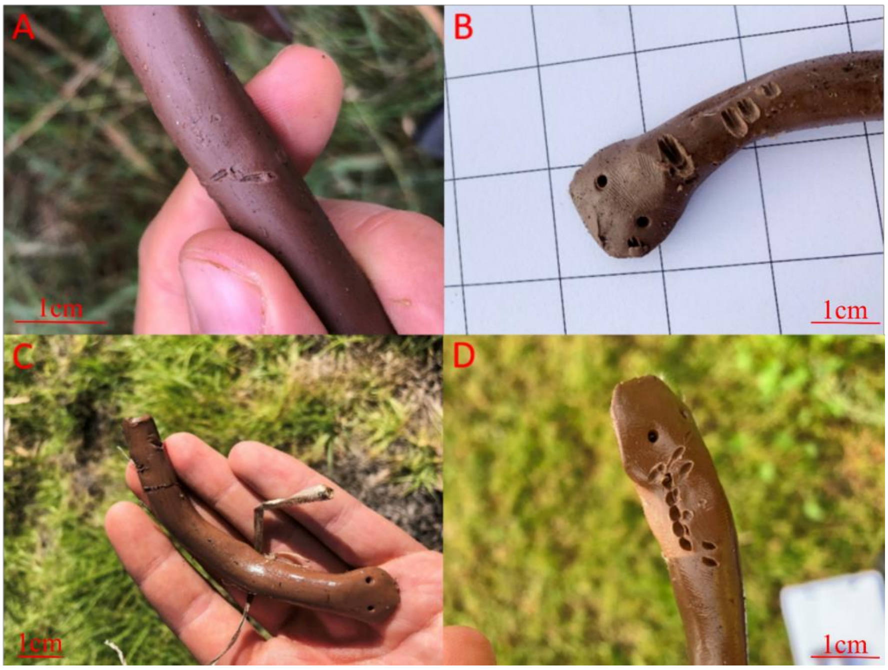


**Fig S2B** Examples of marks left by predators A) a small rodent, classed as exploratory behaviour and omitted from analysis, B) a rodent, C) a bird and D) a cat on artificial juvenile tiger (*Notechis scutatus occidentalis*) snakes deployed in four different wetlands, dominated by either invasive vegetation (Herdsman Lake and Kogolup Lake) or native vegetation (Black Swan Lake and Yanchep National Park) in the Perth region, Western Australia.

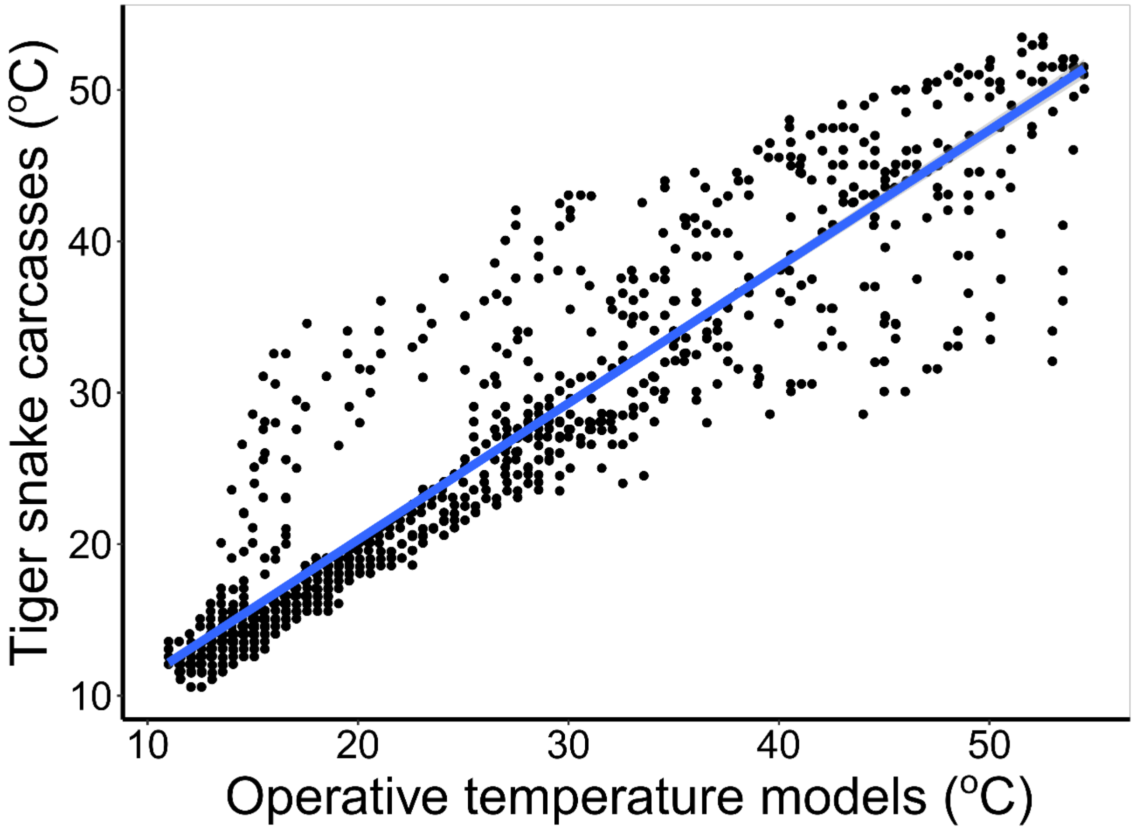


**Fig. S3** Correlation of temperatures between tiger snake (*Notechis scutatus occidentalis*) carcasses and copper pipe operative temperature models. Pearson's correlation: $t_{1149} = 88.2$, $P < 0.001$, slope = 0.90, $R^2$=0.93.

**Results PCA habitat types**

All three PCs differed significantly among habitat types ($F_{5, 234} \leq 149$, $P < 0.001$). For PC1, there was no significant difference between equivalent habitat types in native and introduced grass dominated communities (e.g. KO and NO; *post-hoc* comparisons $P \geq 0.527$); while habitat types in the same plant community (e.g. KO vs KS) did differ (*post-hoc* comparisons $P < 0.001$; Fig 2). For PC2 habitat types within kikuyu grass did not differ (*post-hoc* comparisons $P \geq 0.918$) they did within native vegetation (*post-hoc* comparisons $P \leq 0.007$) except for NO and NW (*post-hoc* comparisons $P = 0.492$). All the habitats within kikuyu grass differed from those in native vegetation (*post-hoc* comparisons $P < 0.001$). PC3 distinguished KS from all other habitat types (*post-hoc* comparisons $P < 0.001$) and KW differed from all the native habitat types (*post-hoc* comparisons $P \leq 0.018$; Fig 2).

**Table S1** Eigenvalues and loading values for the first three principal components (PC; with eigenvalues >1) resulting from principal components analysis of habitat characteristics for western tiger snake (*Notechis scutatus occidentalis*) habitats from four different wetlands, dominated by either invasive vegetation (Herdsman Lake and Kogolop Lake) or native vegetation (Black Swan Lake and Yanchep National Park) in the Perth region, Western Australia.

| Eigenvalues and loading values | PC1 | PC2 | PC3 |
|---|---|---|---|
| Eigenvalues | 2.485 | 1.669 | 1.010 |
| Variation explained (%) | 35.503 | 23.843 | 14.429 |
| Loading values | | | |
| Canopy cover (%) | 0.893 | -0.921 | -0.164 |
| Trees (#) | 0.725 | -0.061 | -0.456 |
| Woody vegetation (#) | 0.577 | -0.341 | 0.081 |
| Ground cover (%) | 0.110 | 0.865 | 0.004 |
| Understory height (cm) | 0.325 | -0.295 | 0.830 |
| Vegetation density (%) | 0.185 | 0.811 | 0.243 |
| Light availability (lux) | -0.821 | -0.212 | -0.140 |

*Operative temperature model calibration*

To determine the paint scheme for the copper models that most closely matched the reflectivity of tiger snakes we conducted several experiments comparing our copper models to tiger snake carcases sourced from a previous study (Lettoof et al. 2020). We painted copper pipes with varying combinations of grey and white, filled them with water and placed an iButton inside each, and then placed them in direct sunlight for 24 hours alongside a tiger snake carcass also containing an iButton. The resulting temperature data were plotted and copper models were compared to the snake carcass by visually inspecting the graph. Once the colour combination that resulted in the closest thermal match was determined, a larger experiment was conducted with four pipes painted grey with two 5cm wide white bands and compared their thermal profile with that of five adult tiger snake carcasses placed outdoors in an exposed location for 48hrs. During this period air temperature ranged from 12.2˚C to 27.2˚C and the snakes and models were exposed to direct sunlight, cloud cover, rain (0.4mm) and variable wind speeds (max 54km/h). Temperature data were recorded at 10min intervals. We calculated thermal properties of the snake carcases and models (mean, minimum, and maximum temperatures, warming and cooling slopes) using a custom written VB program (Visual Basic V6; P.C. Withers). Following this, we conducted a Pearson's correlation test on the body temperatures of tiger snake carcasses and the operative temperatures of the copper models. We then used linear mixed-effects models to compare the temperature characteristics of the snake carcasses and copper models using the various thermal characteristics as dependent variables, treatment (pipes or snake) as a fixed effect and ID (pipe number or carcass number) as a random factor to account for repeated measurements over time. Carcass temperatures were highly correlated with that of the copper models (Pearson's correlation: $t_{1149}$ = 88.2, $P < 0.001$, slope = 0.90, $R^2$=0.93). Further, the mean, minimum, and maximum temperatures, and the warming, and cooling slopes were not significantly different between operative models and snake carcasses ($F_{1, 6} \leq 1.309$, $P \geq 0.296$; Figure S3).

**Table S2** Mean ± SE maximum and minimum operative temperatures ($T_e$) for western tiger snakes (*Notechis scutatus occidentalis*) in each habitat type at sites dominated by kikuyu grass (Herdsman Lake and Kogolup Lake) or native vegetation (Black Swan Lake and Yanchep National Park) over four seasons in the Perth region, Western Australia. Mean ± SE, maximum and the proportion of time the operative temperatures did not deviate from the set-point range (*de* = 0) of tiger snakes in each habitat type in both kikuyu grass and native plant communities (Plant. Com) across four seasons.

| Season | Plant. Com | Habitat | $T_e$ | | | *de* | | |
|---|---|---|---|---|---|---|---|---|
| | | | mean | max | min | mean | max | %=0 |
| Spring | | | | | | | | |
| | Kikuyu | Open | 16.2 ± 0.36 | 33.8 | 4.0 | 7.9 ± 0.53 | 20.0 | 8.3 |
| | | Sedge | 17.0 ± 0.41 | 29.0 | 6.0 | 7.1 ± 0.66 | 17.9 | 9.0 |
| | | Woodland | 18.1 ± 0.42 | 32.0 | 8.5 | 6.1 ± 0.75 | 15.4 | 9.1 |
| | Native | Open | 16.8 ± 0.37 | 34.0 | 6.0 | 7.5 ± 0.73 | 17.9 | 11.6 |
| | | Sedge | 17.2 ± 0.31 | 32.0 | 4.5 | 6.8 ± 0.55 | 19.4 | 5.9 |
| | | Woodland | 17.1 ± 0.39 | 36.8 | 8.5 | 7.1 ± 0.88 | 15.4 | 8.0 |
| Summer | | | | | | | | |
| | Kikuyu | Open | 23.6 ± 0.51 | 38.5 | 10.3 | 3.2 ± 0.84 | 13.6 | 11.2 |
| | | Sedge | 23.3 ± 0.48 | 37.5 | 7.5 | 3.2 ± 0.71 | 16.4 | 9.9 |
| | | Woodland | 22.8 ± 0.48 | 38.0 | 11.5 | 2.7 ± 0.7 | 12.4 | 23.3 |
| | Native | Open | 23.0 ± 0.42 | 42.0 | 7.4 | 3.2 ± 0.79 | 16.5 | 13.7 |
| | | Sedge | 21.8 ± 0.34 | 35.2 | 6.7 | 2.6 ± 0.62 | 14.7 | 17.4 |
| | | Woodland | 22.5 ± 0.44 | 38.5 | 8.5 | 3 ± 0.75 | 14.4 | 12.3 |
| Autumn | | | | | | | | |
| | Kikuyu | Open | 16.9 ± 0.71 | 34.5 | 0.0 | 7.2 ± 0.78 | 25.9 | 10.7 |
| | | Sedge | 16.6 ± 0.79 | 33.5 | 0.5 | 7.5 ± 0.89 | 25.4 | 9.3 |
| | | Woodland | 18.4 ± 0.81 | 36.8 | 1.3 | 6.3 ± 0.96 | 22.6 | 11.4 |
| | Native | Open | 17.3 ± 0.67 | 34.0 | 1.0 | 7.1 ± 0.71 | 22.9 | 17.1 |
| | | Sedge | 17.3 ± 0.66 | 32.5 | 4.0 | 6.9 ± 0.85 | 19.9 | 9.7 |
| | | Woodland | 18.2 ± 0.74 | 38.4 | 5.0 | 6.4 ± 1.07 | 18.9 | 10.7 |
| Winter | | | | | | | | |
| | Kikuyu | Open | 11.0 ± 0.37 | 22.0 | 0.0 | 12.9 ± 0.33 | 24.0 | 0.0 |
| | | Sedge | 11.4 ± 0.37 | 24.5 | 0.0 | 12.5 ± 0.4 | 23.9 | 0.0 |
| | | Woodland | 12.7 ± 0.35 | 23.0 | 3.0 | 11.2 ± 0.41 | 20.9 | 0.0 |
| | Native | Open | 12.1 ± 0.34 | 23.5 | 1.5 | 11.8 ± 0.5 | 22.4 | 0.0 |
| | | Sedge | 12.6 ± 0.31 | 22.0 | 3.0 | 11.3 ± 0.42 | 20.9 | 0.0 |
| | | Woodland | 12.4 ± 0.32 | 19.0 | 3.5 | 11.5 ± 0.28 | 20.4 | 0.0 |
